# Phytochemical and antibacterial analyses of *Onychium japonicum* (Thunb.) Kunze

**DOI:** 10.1101/2022.06.14.496165

**Authors:** Syed AHsan Shah, Ghulam Mujtaba Shah, Mohamed M. Hassan, Walaa F. Alsanie, Shamyla Nawazish, Waqar Ali, Abdul Basit, DR. Nosheen Shafqath, Nadia Jabeen, Syed Abdul Majid Shah, Zarbad Shah, Muhammad Ishtiaq, DR. Alia Gul, jan alam, Muhammad Islam, Asma Faheem, Experts from Microbiology and Biichemistry as well participated in the current work

## Abstract

In clinical practice bacterial resistance against antibiotics has become a serious health problem, thus using alternative approaches such as natural products as the supplementary drug could solve it. Therefore, the current study was conducted to thoroughly investigate the enrichment of phytochemicals and antibacterial potential of *O. japonicum*. The fronds samples of *O. japonicum* prepared in different solvents were used against MDR bacterial strains and phytochemical analyses.

The analyses of data revealed that *O. japonicum* was enriched with flavonoids, alkaloids, saponins, tannins, glycosides, carotenoids, terpenoids, phlobatanins, phenols, and coumarins while quantitatively this plant has a significantly higher content of phenols(1100.91µM/g) followed by sugar contents (748.67µM/ml), ascorbic acid (426.12mM/g), flavonoids (160.65mg/g), anthocyanin (101.06µM/g) and proline (80.58µM/g). On the other hand, the organic extracts of *O. japonicum* were highly active against all bacterial strains while hydric extract was inactive against selected bacteria. Specifically, *O. japonicum* was highly active against *S. aureus* in all organic extracts (chloroform=16.66±0.33, ethyl acetate=15±0.57, methyl alcohol= 14±0.57, N-hexane=20.33±0.33) followed by *K. pneumonia* (chloroform=14.33±0.33, ethyl acetate=4.33±4.33, methyl alcohol=3.66±3.66, N-hexane=16.66±0.33) and *P. aeruginosa* (chloroform= 8.33±4.17, ethyl acetate=8.33±4.17, methyl alcohol=6±3.00, N-hexane=9.33±4.66), while *E. coli* (chloroform=0±0.00, ethyl acetate=7±3.51, methyl alcohol=3.33±3.33, N-hexane=4±4.00).

Based on current findings it is concluded that *O. japonicum* is enriched in many useful phytochemicals that could be use as a supplement with other traditionally used antibiotics.

## 1. Introduction

Extensive use of antibiotics made bacteria more intelligent thus they develop new pathways to resist many drugs (Fankam et al., 2017). To overcome multidrug drug resistance, clinician usually uses a combination of drugs (Ahmad et al., 2014), or add additional natural products as supplements (Asher et al., 2017). Prolonged use of drugs also causes some serious adverse effects on the patient’s health such as kidney failure or liver toxicity (Andrad & Tulkens, 2011). However, medicines are obtained from natural sources or using a natural source (medicinal plant) directly as medicine has comparatively fewer toxic effects (Veeresham, 2012).The plants are the basic source of natural products and used as traditional medicines since the history of human beings, that approximately goes back to 60,000 years, which was estimated after the discovery of 5000 years old Sumerian clay slabs used for the preparation of the plant drugs (Akhtar & Foyzun, 2017). In recent years, the use of plant-driven medicines has gotten much attention due to their high benefits and fewer side effects on human health (Aye et al., 2019). Such as *Viola odorata* is being used as an antidepressant (Feyzabadi et al., 2014) and compounds taken from *Cannabis sativa* are commonly prescribed for muscle strengthening in Epilepsy, dystonia, and Parkinson’s disease (Freeman et al., 2019). Although synthetic organic chemistry has significantly grown in the 20^th^ century, around 25% of the standard drugs are being directly or indirectly obtained from plants (Baskaran et al., 2018). Most importantly, the plants have high medicinal power due to the enrichment of various secondary metabolites (Pertibha et al., 2014). In recent years a significant rise in the unprescribed use of antibiotics and other drugs has been observed which caused serious health problems and various bacterial strains have developed resistance to multiple drugs (WHO, 2020). Therefore, the demand for new antibacterial agents has significantly increased, and medicinal plants for their safe and recyclable bioactive chemical constituents have also been prioritized (Rakkimuthu et al., 2018).

Pteridophytes have been known for more than 2000 years for their use in folk medicines. Medicinal use of pteridophytes is also recommended by Ayurvedic systems of medicine and the Unani system of medicines (Panda et al., 2014). Notably, pteridophytes play a vital role in folklore medicines and are a vital source of food that can be used in the prevention of diseases and for the maintenance of human health (Ismail et al., 2014). Ethyl acetate friction of *Azolla microphylla Kaulf* an aquatic pteridophytes, resist the growth of pathogenic bacterium X. oryzae (Abraham et al., 2015). Methanolic extract of eleven ptridophytic species were tested against *s. typhi, v. cholerae* and *p. aeruginosa*. Methanolic extract of selected pteridophytes show best activity against selected bacteria (Maridass, 2021).

Our experimental plant, *O. japonicum* belong to family Pteridaceae related to this medicinally potent group of plants (Pteridophyte). Chalcone derivative extracted from *O. japonicum* showed high cytotoxic activity against human cervical cancer and hepatocellular carcinoma cell lines Hela cells and BEL-7402 cells, respectively (Li et al., 2007). Two new compounds, cyathane diterpene glucosides (1 and 2) were isolated from *O. japonicum* were found to have anti-inflammatory effects (Ke Zan et al., 2021). The methanolic extracts of *O. japonicum* contain significantly high insecticidal activity against adult house flies and mosquitoes (Huang et al., 2010). Similarly, the petroleum ether extracts *O. japonicum* was more active against gram-positive bacteria as compared to gram-negative bacteria and fungi (Khan & Shaheen, 1998).

There is phytochemical exploration and antibacterial studies present in literature on *O. Japonicum* but the current study was conducted to explore a broad picture of its phytoconstituents and antibacterial potential in different solvents against various multidrug resistant bacterial strains.

## 2. Materials and methods

### 2.1. Plant collection and processing

Fern *O. japonicum* was collected from District Mansehra and processed at the Department of Botany Hazara University Mansehra, KPK, Pakistan. First of all, fronds of the plant were air-dried and powdered by using a commonly used blender. 50g powdered samples were dissolved in 400ml of 5 different solvents (Chloroform, Methyl alcohol, N-hexane, Ethyl acetate, and Distilled water) respectively and kept in a shaker for 7 days at room temperature (25±2°C). Then each sample was filtered out using filter paper (F1001 grade) and subjected to a rotary evaporator for completely drying and concentrating samples, and stored at 4°C till further analyses. All the sampling and experimental procedures were approved by the advance studies & research board (ASRB) and ethical committee of Hazara University, Mansehra, Pakistan.

### 2.2. Phytochemical analyses

Plant samples were prepared in different solvents for determining various phytochemicals; different phytochemicals were detected based on changes in the solvents’ colors. For terpenoids, flavonoids, saponins, tannins, phenols, carbohydrates, and glycosides we followed the protocols published by (Iqbal et al., 2019), while alkaloids, Phlobatanins, coumarins, and carotenoids were analyzed by standard protocols mentioned in previous studies (Ali et al., 2020). To quantify the phytochemicals in *O. japonicum*, we used a spectrophotometer (UV 1900), by following the manufacturer’s guidelines. Specifically, flavonoids and phenols (Sawhey and Jassal 2013), sugar content (Pawar and Mello, 2011), anthocyanin (Lao et al., 2016), ascorbic acid (senguttuvan et al., 2014), and prolines (Cariilo and Gibon, 2011) were accordingly determined by following the modifications mentioned in the studies.

### 2.3. Antibacterial activity

Four bacterial MDR strains (*Pseudomonas aeruginosa, Escherichia coli, Staphylococcus aureus*, and *Klebsiella pneumonia*) were used to determine the antibacterial potential of *O. japonicum*. Dry extracts were dissolved in DMSO (Dimethyl Sulphoxide) in a proportion of 120mg/ml and the disc diffusion method was applied to determine the antibacterial effects of plant samples. Standard antibiotic levofloxacin (5µg) and DMSO were used as positive and negative control/blank respectively.

### 2.4. Statistics

Statistical analyses were performed by using software (statistix 8.1). All experiments were performed in triplicate, and the mean of replicates was noted. The ANOVA test was performed to observe significant differences among different treatments. A P-value ≤ 0.05 was considered significant.

## 3. Results

### 3.1. Phytochemical screening

### 3.2. Qualitative phytochemical analysis

The qualitative analysis showed that *O. Japonicum* enriched with different compounds such as flavonoids, carotenoids, terpenoids, Phlobatanins, alkaloids, saponins, carbohydrates, glycosides, phenols, tannins, and coumarins (Table 1). We used different solvents to obtain a better picture of different compounds presence in *O. Japonicum*, thus some variations were found in different detection methods. Different compounds produced different colors in different solvents (Figure 1A-E). In short, the flavonoids were strongly detected in methyl alcohol, N-hexane, and distilled water extracts but did not observe in chloroform and ethyl acetate extracts. While, the carotenoids were strongly observed in chloroform, ethyl acetate, and N-hexane extracts but have not been detected in the methyl alcohol and distilled water extracts. On the other hand, the terpenoids showed high concentration in methyl alcohol and distilled water, while the moderate concentration in chloroform and N-hexane extracts, and absent in extracts made in ethyl acetate. Similarly, the concentration of phlobatanins was moderately detected in chloroform, ethyl acetate, and methyl alcohol extracts and was not seen in the N-hexane and distilled water extracts. The alkaloids showed their strong presence in chloroform and ethyl acetate extracts, moderately detected in N-hexane and distilled water extracts while no traces of alkaloids were detected in samples prepared in the methyl alcohol. Saponins were moderately present in the methyl alcohol extract, while absent in all other extracts. Carbohydrates were absent in distilled water, ethyl acetate, and N-hexane extracts while present in methyl alcohol and chloroform extract. Moderate presence of glycosides was observed in chloroform, methyl alcohol, and distilled water extracts while absent from other extracts. Phenols were strongly present in chloroform and methyl alcohol extracts, while absent from all other extracts. The same results were observed for tannins. Distilled water and methyl alcohol extracts showed a high concentration of coumarins and were moderately detected in N-hexane and ethyl acetate extracts, while absent from chloroform extract. These findings show that methyl alcohol and chloroform solvents are preferable for detecting most of the phytochemicals.

**Table 1.** Phytochemical constituents of *Onychium japonicum* in different extracts.

| Tests | Chloroform | Ethyl acetate | Methyl alcohol | N-hexane | Distilled water |
| --- | --- | --- | --- | --- | --- |
| **Flavonoids** | - | - | +++ | +++ | +++ |
| **Carotenoids** | +++ | +++ | - | +++ | - |
| **Terpenoids** | ++ | - | +++ | ++ | +++ |
| **Phlobatanins** | ++ | ++ | ++ | - | - |
| **Alkaloids** | +++ | +++ | - | ++ | ++ |
| **Saponins** | - | - | ++ | - | - |
| **Carbohydrates** | +++ | - | ++ | - | - |
| **Glycosides** | ++ | - | ++ | - | ++ |
| **Phenols** | +++ | - | +++ | - | - |
| **Tannins** | +++ | - | +++ | - | - |
| **Coumarins** | - | ++ | +++ | ++ | +++ |

**Figure 1.**
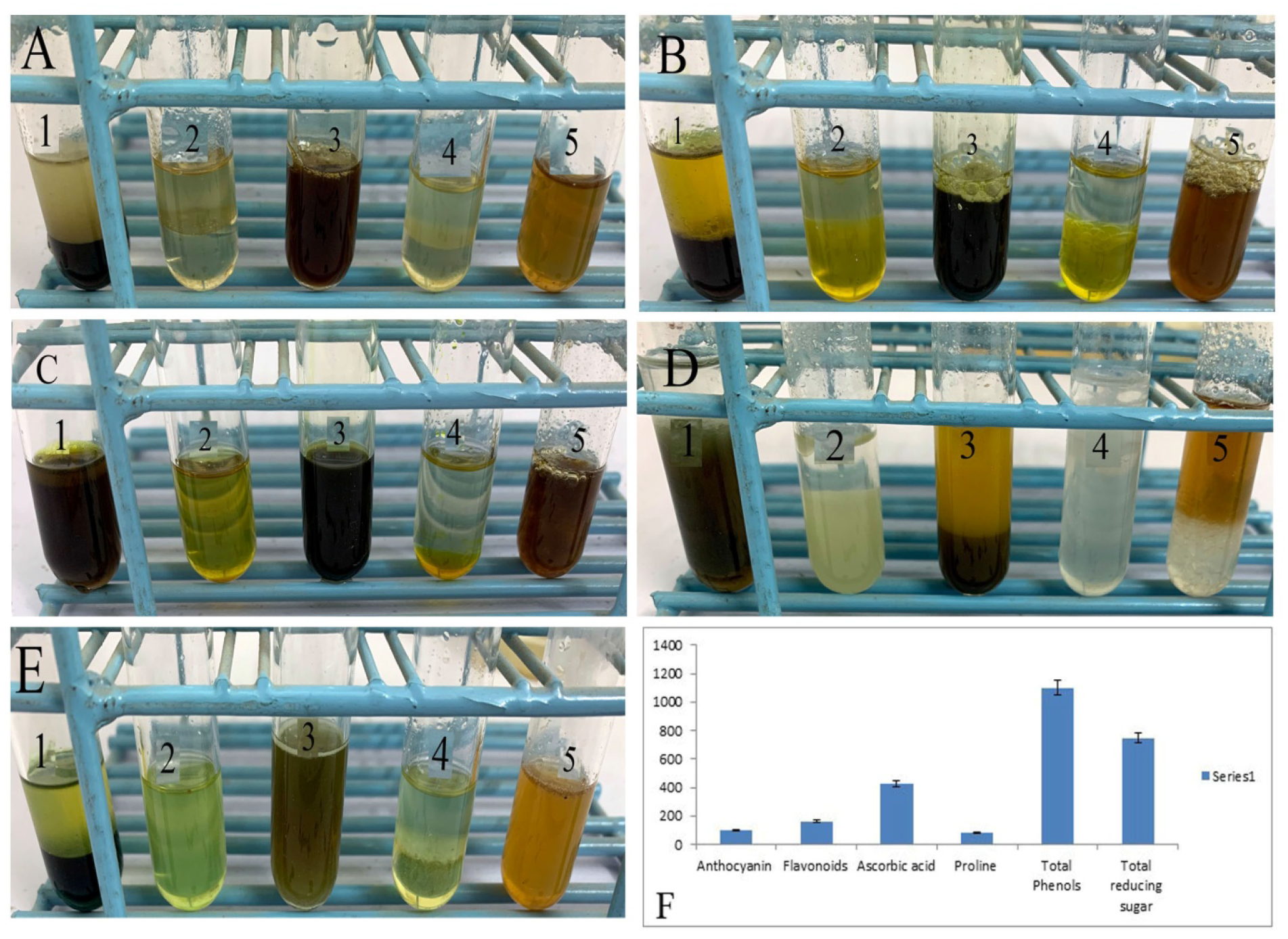
Phytochemical analysis of *O. japonicum*. (A) shows the test for coumarins, (B) tannins, (C) phenols, (D) glycosides, (f) alkaloids, and (F) graphical representation of quantitative phytochemicals. **Labels on tubes** (1= chloroform extract, 2= ethyl acetate extract, 3= methyl alcohol extract, 4= N-hexane extract, 5= distilled water extract).

### 3.3. Quantitative analysis of phytochemicals

Furthermore, we quantified some of the phytochemicals detected in the *O. Japonicum*. Analysis of the data showed that *O. japonicum* has a significantly high amount of phenols (1100.91µM/g), followed by total sugar content (748.67 µM/ml). Similarly, we also measured the concentration of ascorbic acid (426.12 mM/g), flavonoids (160.65 mg/g), anthocyanin (101.06 µM/g), and proline (80.58 µM/g) in the *O. japonicum* (Figure, 1F).

### 3.4. Antibacterial activity

Through phytochemical analyses, we assumed that *O. japonicum* may have strong antibacterial potential thus we analyzed this plant against multidrug-resistant strains of bacteria. As mentioned earlier that we prepared samples in different solvents like distilled water, chloroform, methyl alcohol, ethyl acetate, and N-hexane. Using standard bacterial culturing and drug screening procedures we applied different concentrations of the plant samples on bacteria. Overall, we found that *O. japonicum* showed significant antibacterial potential against tested bacteria in all extracts except distilled water extract (Figure 2). Particularly, *O. japonicum* was found to have the best bactericidal activity against *S. aureus* (Figure 2D), as compared to other bacteria. The N-hexane extracts of the plant showed maximum activity (20.33±0.33) against *S. aureus* as compared with chloroform extracts (16.66±0.33), ethyl acetate (15±0.57), and methyl alcohol (14±0.57). Similarly, N-hexane extracts also showed excellent activity (20.33±0.33) against *K. pneumonia* followed by chloroform (14.33±0.33), ethyl acetate (4.33±4.33), and methyl alcohol (3.66±3.66) extracts. The extracts of *O. japonicum* were observed as less active against *P. aeruginosa* and *E. coli*, the maximum activity showed by N-hexane extract (9.33±4.66), followed by chloroform (8.33±4.17) and ethyl acetate (8.33±4.17) and methyl alcohol (6±3.00) extracts against *P. aeruginosa* while against *E. coli* ethyl acetate, N-hexane and methyl alcohol showed (7±3.51), (4±4.00) and (3.33±3.33) activity, respectively, however, the extracts prepared in chloroform were inactive against *E. coli*. The standard antibiotic levofloxacin (5µg) disc was used as positive control while DMSO was used as a negative control or blank of the study (Table 3).

**Table 2.** Quantitative analysis of *Onychium japonicum*.

| Phytochemicals | Percentage ± standard error |
| --- | --- |
| Anthocyanin ($\mu\text{M/g}$) | 101.06±5.45 |
| Flavonoids (mg/g) | 160.65±13.97 |
| Ascorbic Acid (mM/g) | 426.12±4.27 |
| Proline ($\mu\text{M/g}$) | 80.58±1.195 |
| Phenols (mg/ml) | 1100.91±43.46 |
| reducing sugars<br>($\mu\text{M/ml}$) | 748.67±9.96 |

**Table 3.** Antibacterial activity of *Onychium japonicum*.

| S.NO | Strains | CH | EA | MA | NH | DW | LEVO | DMSO |
| --- | --- | --- | --- | --- | --- | --- | --- | --- |
| 1 | *Staphylococcus aureus* | 16.66±0.33 | 15±0.57 | 14±0.57 | 20.33±0.33 | 00 | 24.66±0.88 | 00 |
| 2 | *Klebsela pneumonia* | 14.33±0.33 | 4.33±4.33 | 3.66±3.66 | 16.66±0.33 | 00 | 14±1.15 | 00 |
| 3 | *Escherichia coli* | 0±0.00 | 7±3.51 | 3.33±3.33 | 4±4.00 | 00 | 20.33±0.88 | 00 |
| 4 | *Pseudomonas aeruginosa* | 8.33±4.17 | 8.33±4.17 | 6±3.00 | 9.33±4.66 | 00 | 19±0.57 | 00 |

**Zone of inhibition mm ± Standard Error means**
**Key- CH= chloroform, EA= ethyl acetate, MA=methyl alcohol, NH= n-hexane, DW=distilled water, LEVO= Levofloxacin, DMSO = dimethyl sulphoxide.**

**Figure 2.**
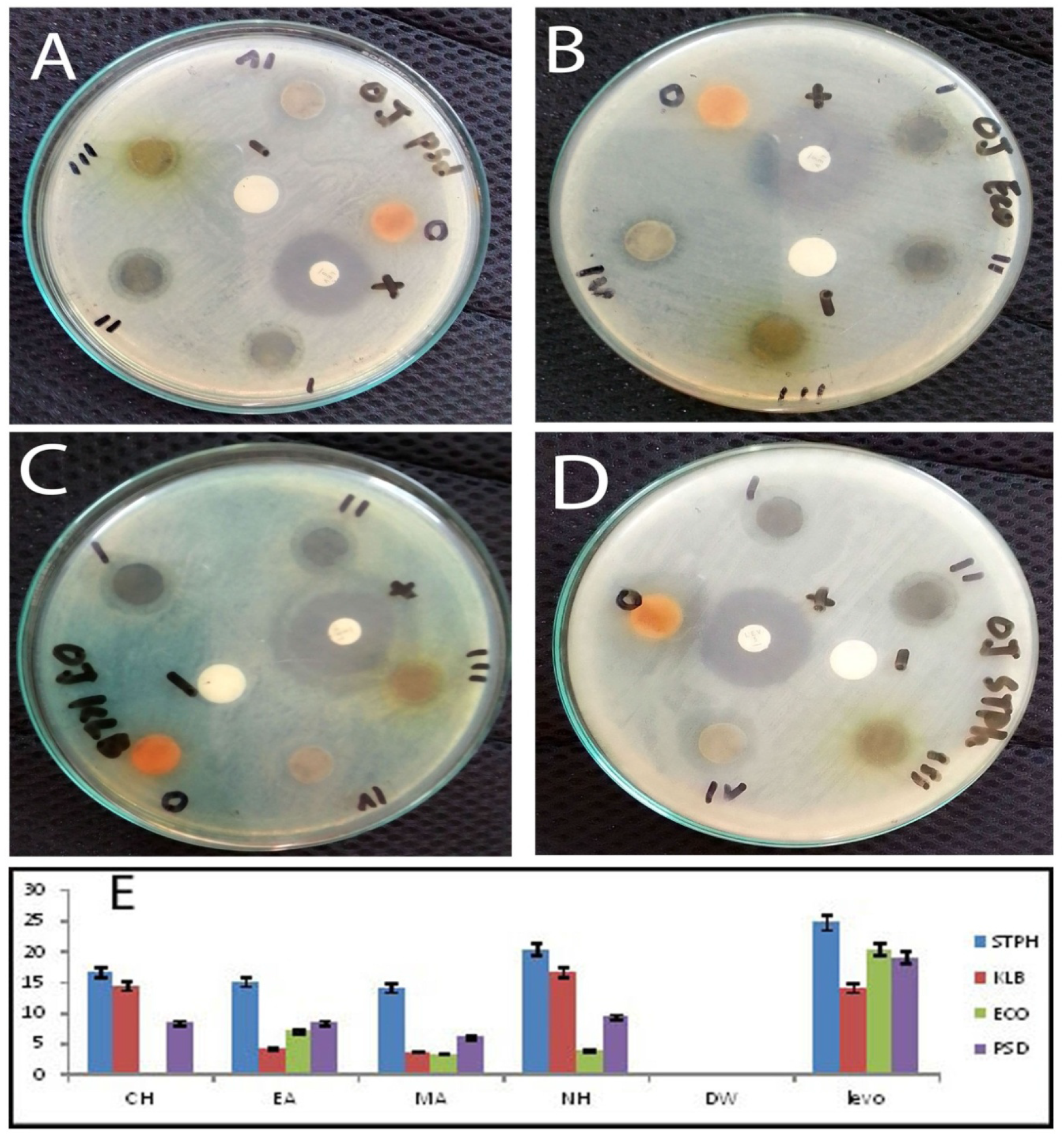
Antibacterial activity of *O. japonicum* five extracts (I = chloroform extract, II= ethyl acetate extract, III= methyl alcohol extract, IV= N-hexane extract, 0= distilled water extract, + = positive control, - = negative control). **against** (A) *P. aeurogenosa*, (B) *E. coli*, (C) *K. pneumonia*, (D) *S. aureus*, and (E) graphical representation of the antibacterial activity of *O. japonicum*.

## 4. Discussion

The clinical use of natural products has gotten much attention in recent years due to their better clinical outcomes and fewer adverse effects (Yuan et al., 2016). Medicinal plants (NPs) are regarded as the best source for drug discovery due to their enrichment in a variety of chemicals (Zhang et al., 2020) like alkaloids, terpenes, glucosinolates, phenolic compounds, flavonoids, and metabolites (Agyare et al., 2019). Aspirin, a blood thinner drug extracted from *Filipendula ulmaria* (Anand et al., 2019), *Digitalis purpurea L*. produces tiotropium bromide a drug used for respiratory diseases (Mundy & Kirkpatrick, 2004). Similarly, *O. japonicum* is a very important medicinal fern, showing anticancer activity (Li et al., 2007), anti-inflammatory (Ke Zan et al., 2021), and antibacterial activity (Khan & Shaheen, 1998). In the current study, we provide a broad picture of its biochemicals, and also illustrated the efficacy of different methods. We used five solvents to identify the phytochemicals and found samples prepared in methyl alcohol and chloroform showed good efficacy for the identification of phytochemicals, specifically, higher phytoconstituents were detected in the methyl alcohol followed by chloroform extract. Similar findings have been reported by (Chigayo et al., 2016), claiming that methyl alcohol is the best solvent for the detection of different classes of phytochemicals. Particularly, chloroform and methyl alcohol are the best solvents for identifying alkaloids, tannins, sugar, saponins, flavonoids, terpenoids, cardiac glycosides, phenolics, and anthraquinone (Ali et al., 2020). In addition, N-hexane has also been considered the best solvent for analyzing different classes of phytochemicals (Ramesh, 2016). We successfully identified flavonoids, alkaloids, saponins, tannins, glycosides, carotenoids, terpenoids, phlobatanins, phenols, and coumarins compounds in *O. japonicum*. Individually these compounds have medicinal and commercial importance (Mujeeb et al., 2014).

It has been confirmed that secondary metabolites contribute to the antimicrobial potential of the medicinal plants by different molecular mechanisms such as tannins inhibit protein synthesis by forming permanent inhibitory complexes with proline-rich proteins. Clinically, tannins have been used against inflammation and injuries (Pizzi, 2021). Herbs containing tannins as a major constituent have been suggested to treat intestinal disorders like diarrhea and dysentery (Fraga et el., 2021).

We believe that the antibacterial activity of the *O. japonicum* is due to the presence of enormous amount of biologically active phytochemicals. In *O. japonicum* we observed a higher concentration of alkaloids which are important secondary metabolites potentially cytotoxic compounds thus *O. japonicum* possesses high anticancer activity (Nobori et al., 1994) and also have powerful analgesic effects (pietta PG, 2000). Saponins show anti-inflammatory effects (González et al., 2020). Flavonoids are well-known antimicrobial, analgesic, antioxidant, anti-allergic, cytostatic anti-inflammatory compounds (Ullah et al., 2020). Phlobatanins revealed anti-inflammatory, analgesic and wound-healing properties and are good to use in cardiovascular disorders due to their antioxidant character (Emmanuel et al., 2020). Similarly, carotenoids exhibit antioxidant activity; in addition, they are also responsible for a pro-vitamin A function, helping to control age-related eye disorders (Eggersdorfer and Myss, 2018). Yang et al., 2020 reported antimalarial, antiviral, antibacterial, and anti-inflammatory activity for terpenoids. Interestingly, different drugs are glycosides in nature like cardiac glycosides and some antibiotics like streptomycin (Britannica, 2020), glycosides can be used to treat heart disorders also use as purgative, antioxidant, and anti-inflammatory (Britannica, 2018). Similarly, phenols have antioxidants and free radical scavenging properties which help protect against different cardiovascular diseases, inflammation, bacterial infections, and several neurodegenerative diseases (Ozcan et al., 2014). Coumarins are used to treat prostate cancer, renal cell carcinoma, leukemia, and also to cope with the side effects of photochemotherapy (Kupeli et al., 2020). In the current study, we detected plenty of phytochemicals in *O. japonicum*, which reveals that this plant has high medicinal power that is yet to be explored. We believe it could be used in several diseases specifically cancer, cardiovascular diseases, other inflammatory diseases, and bacterial infections.

To further explore the antibacterial effects of *O. japonicum* we specifically choose multidrug-resistant bacterial strains. The extracts prepared in the organic solvents showed high antibacterial activity while hydric extract was inactive against all tested bacteria. We speculate that all active chemicals or compounds of the *O. japonicum* might be non-polar in nature thus could not be dissolved in water. Previously, the *O. japonicum* has shown strong activity against gram-negative bacteria (Khan & Shaheen, 1998). Furthermore, *O. japonicum* was highly active against *S. aureus*, and *Pseudomonas aeruginosa*, while least active against the *E. coli* and *K. pneumonia* showing that this plant could potentially be used against the infections caused by *S. aureus*, and *P. aeruginosa*. Particularly, *S. aureus* is considered as one of the most sensitive bacterial strains that could easily be killed by various natural or synthetic drugs (Thilza et al., 2010; Farjan et al., 2014). On the contrary side, *E. coli* and *K. pneumonia* have been considered as the most resistant bacterial strains which require high potency of drugs or combination therapy for complete eradication (Muchtaromah et al., 2020; Mostafa et al., 2018). Based on current findings we believe that *O. japonicum* is an important medicinal plant, to elaborate its clinical importance extensive studies involving cell lines and animal models are required. Our findings have some limitations; although we detected several important phytochemicals in the *O. japonicum* due to the lack of advanced sophisticated techniques many vital chemicals and heavy metals might have not been detected. We were unable to investigate anticancer effects on many cancer cells lines or xenograft mice and anti-inflammatory effects or other important clinical aspects. The antibacterial studies should also be expanded from Petri-plate research to clinical studies in animals leading to human clinical trials.

## 5. Conclusion

In summary, we thoroughly investigated *O. japonicum* for the presence of biologically active phytochemicals and antibacterial potential against multidrug-resistant bacterial strains. Several phytochemicals including flavonoids, alkaloids, saponins, tannins, glycosides, carotenoids, terpenoids, phlobatanins, phenols, and coumarins were detected and while phenols, sugar contents, ascorbic acid were found to have higher enrichment in *O. japonicum*. Moreover, the organic extracts of *O. japonicum* were found to have high activity against all tested bacteria while hydric extracts did not show any bactericidal activity. In short, *S. aureus* was the most sensitive strain while *E. coli* was detected as the most resisting bacterial strain against *O. japonicum*. Based on these findings, we conclude that *O. japonicum* has many important clinical functions including antibacterial, which require further clinical validation.

## 6. Acknowledgments

The authors extend their appreciation to Taif University for funding current work by Taif University Researcher Supporting Project number (TURSP-2020/53), Taif University, Taif Saudi Arabia.

We also acknowledge the laboratotories staff of Department of Botany, Microbiology and Biochemistry of Hazara University. The current study was funded by the IFS project (1-3-F-6135-1).

## 7. Conflict of interest

All authors declare that they have no conflict of interest.

## 8. Data availability

Data generated in such studies such as raw data and figures could be asked from corresponding author on reasonable request.

## Qualitative phytochemical analysis

## Quantitative analysis of phytochemicals

## Notes

### Competing Interest Statement

The authors have declared no competing interest.

